# Drought sensitivity of leaflet growth, biomass accumulation, and resource partitioning predicts yield in common bean

**DOI:** 10.1101/736199

**Authors:** Amber Hageman, Milan O. Urban, Elizabeth Van Volkenburgh

## Abstract

While drought limits yield largely by its impact on photosynthesis and therefore biomass accumulation, biomass is not the strongest predictor of yield under drought. Instead, resource partitioning efficiency, measured by how much total pod weight is contained in seeds at maturity (Pod Harvest Index), is the stronger correlate in *Phaseolus vulgaris*. Using 20 field-grown genotypes, we expanded on this finding by pairing yield and resource partitioning data with growth rates of leaflets and pods. We hypothesized that genotypes which decreased partitioning and yield most under drought would also have strongest decreases in growth rates. We found that while neither leaflet nor pod growth rates correlated with seed yield or partitioning, impacts to leaflet growth rates under drought correlate with impacts to yield and partitioning. As expected, biomass production correlated with yield, yet correlations between the decreases to these two traits under drought were even stronger. This suggests that while biomass contributes to yield, biomass sensitivity to drought is a stronger predictor. Lastly, under drought, genotypes may achieve similar canopy biomass yet different yields, which can be explained by higher or lower partitioning efficiencies. Our findings suggest that inherent sensitivity to drought may be used as a predictor of yield.

**HIGHLIGHT:** In common bean, higher biomass accumulation under drought alone does not guarantee higher yield, as maintenance of higher growth rates and partitioning processes act as an additional requirement.

## INTRODUCTION

Among abiotic stresses, drought has the most detrimental impact on seed yield in the common bean (*Phaseolus vulgaris* L.) (Thung and Rao, 1999). While drought limits yield largely by its impact on photosynthesis and therefore canopy biomass accumulation, findings show that increasing canopy biomass alone does not necessarily lead to an increase in seed yield (Shibles and Weber, 1996). Instead, the ability to partition resources efficiently towards reproductive structures appears to provide the stronger increase to seed yield under drought (Omae *et al.*, 2012; Polania *et al.*, 2016). In particular, one important correlate is Pod Harvest Index (PHI), a measure of photosynthate mobilization from pods (fruits) into seeds (Assefa *et al.*, 2013). Plants with higher PHI values partition greater amounts of pod biomass into seeds, increasing seed yield and resource use efficiency. This means that while some plants can amass a large amount of canopy biomass under drought, only the ones most efficient at moving, or allocating, those resources into their seeds obtain high yield. This raises the simple yet unanswered question: what makes genotypes differ in their ability to allocate resources toward seed production under drought?

Organs that accumulate resources from the rest of the plant (sinks) must obtain resources from other organs that produce or store those resources (sources). Since drought strongly hinders photosynthesis, this is likely to impair source availability. If a plant is source-limited, typically due to photosynthetic limitations, there may be too few resources available to allocate to sinks. However, studies show that seed filling in *Phaseolus vulgaris* is at most only partially coupled with photosynthesis (Smith, 2017). When source resources are not limiting, the rate at which they flow between sources and sinks is largely determined by active allocation processes, such as phloem loading, sugar metabolism, and the sink’s ability to take up and utilize these resources (Farooq *et al.*, 2009). These processes are controlled by signaling as opposed to substrate availability, although sucrose itself acts as a signal regulating many of these processes (Liu, Offler, and Ruan 2013). Different sink organs, or the same organ within different genotypes, can vary widely in the rate at which they metabolize and take up resources, impacting that organ’s ability to attract photosynthate to be delivered to it. For example, *Phaseolus vulgaris* seeds among different genotypes have been shown to take up sucrose at differing rates, even when there were no differences in available sucrose in the sap surrounding the seed (Tegeder *et al.*, 2000) and their growth rates have been shown to correlate with activity of enzymes involved in carbohydrate metabolism, such as invertase and sucrose synthase (Wardlaw, 1990). Since seed production in *Phaseolus vulgaris* is hindered by drought, even in genotypes with high canopy biomass, we predicted that susceptible genotypes may be impacted by a weakening of their ability to take up resources – quantified as sink strength. If true, perhaps whatever signal/s limit allocation, uptake, or use of resources in seeds may also affect these same processes in growing leaves or pods.

A greenhouse experiment showed good correlation between leaf growth rate and seed yield in *Phaseolus vulgaris* lines (Banan and Van Volkenburgh, 2012). While growth rates are not necessarily a measure of resource acquisition, especially under water deficit (Muller *et al.*, 2011), we hypothesized that they may act as an approximation for sink strength and drought sensitivity, such that genotypes whose leaves or pods maintain high growth rates under drought may also achieve higher seed yields and PHI values. In this study, our objective was to test this hypothesis by comparing leaflet growth rates (LGR), pod growth rates (PGR), seed yield, and resource partitioning efficiency (via PHI) to one another and determine drought’s impacts on these processes. In this study, the ‘impact’ on a trait specifically refers to the percent decrease in value between well-watered and droughted plants, where genotypes with larger decreases are considered more impacted. While many studies have previously compared various lines’ agronomy and phenology in the field (e.g., Beebe et al. 2013; Polania et al. 2016; I. M. Rao et al. 2017; Smith 2017), data on growth rates of pods and leaves collected alongside these measurements, with quantifications of the impacts to these traits under drought, are lacking. This study aimed to fill that gap to: 1) understand whether impacts on growth rates and partitioning efficiency under drought relate to drought resistance (high seed yield) and 2) look at impacts on growth under drought in different tissue types to better understand systemic drought responses.

## MATERIALS & METHODS

### Plant material

We conducted a field study using 19 lines of common bean (*P. vulgaris* L.) and one line of tepary bean (*P. acutifolius*). These 20 genotypes were chosen to represent a wide range of observed PHI values in field grown plants (Table 1), providing variability for probing physiological responses. Sixteen of these genotypes were made up of two RIL (recombinant inbred lines) populations, including the four parents and 6 RILs from each cross. These RIL populations were created using parents (MD23-24 x SEA5 – MR RIL) and (BAT881 x G21212 – BH RIL) which differed in their response to abiotic stress, such that their offspring would mostly fall between them in traits related to stress resistance, including PHI (Polania *et al.*, 2017; Diaz *et al.*, 2018). The remaining lines, SEN56, INB841, and DOR390, were chosen as routine checks; DOR390 for drought-sensitivity, INB841 for drought resistance, and SEN56 for high pod partitioning efficiency. *P. acutifolius* (G40001) was included since it is highly drought tolerant and has very high PHI.

**Table 1.**
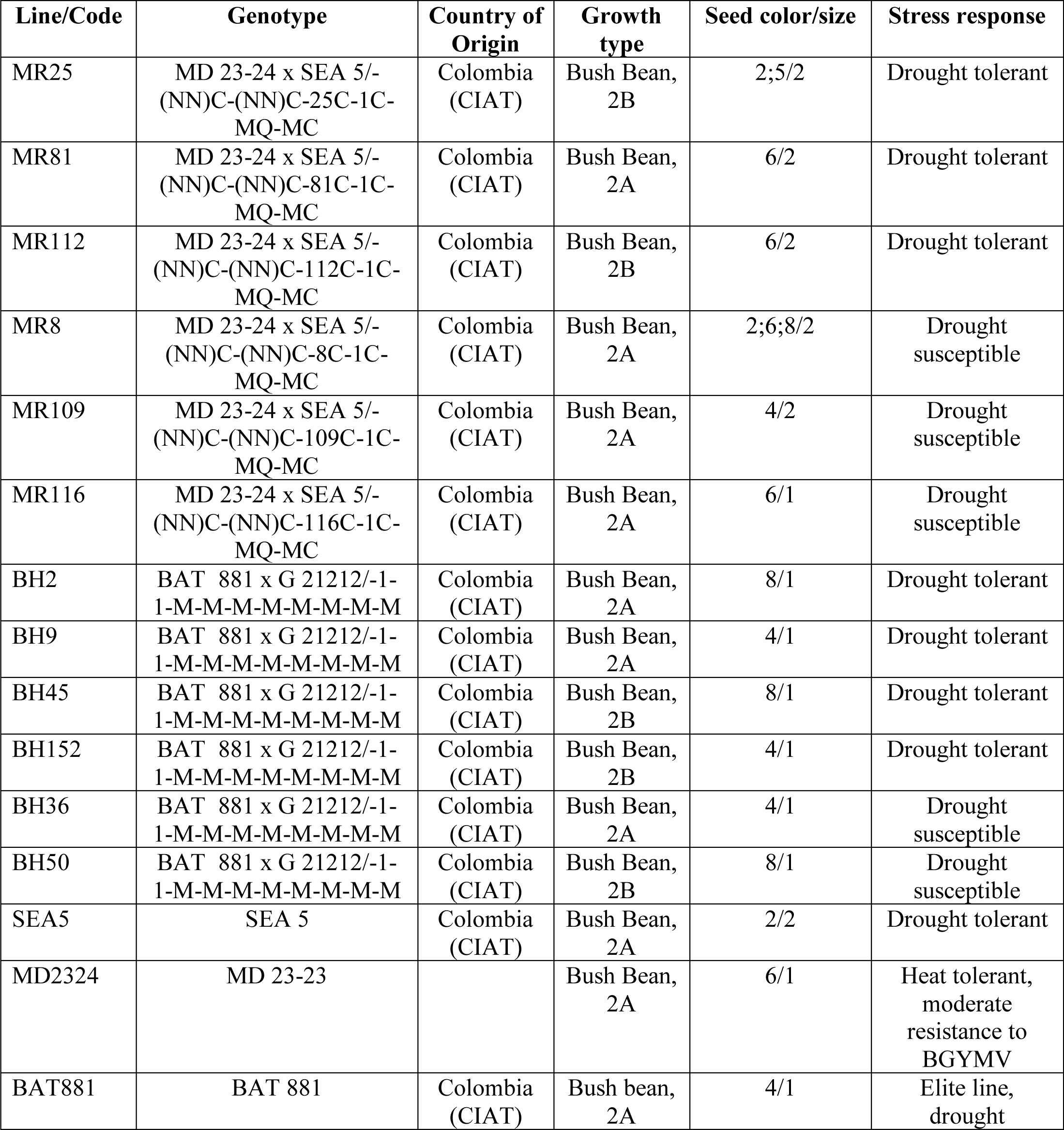
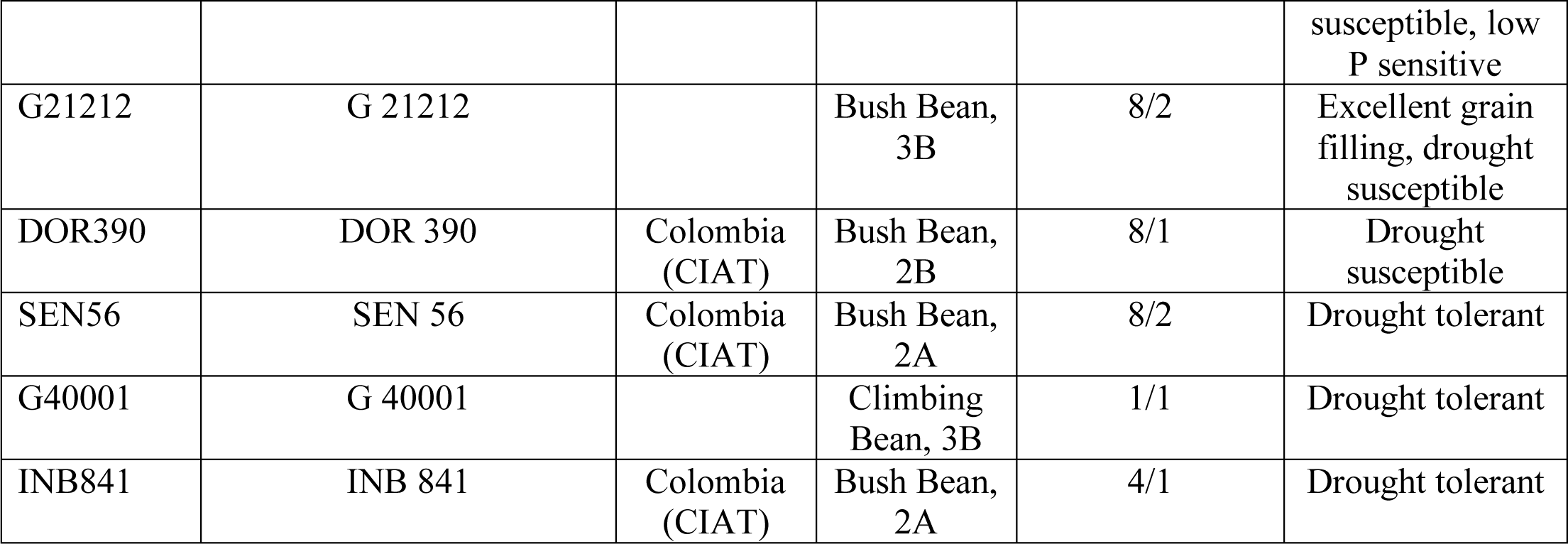
Growth type (2A = indeterminate bush habit, erect stems without guide; 2B = indeterminate bush habit, erect stems with guide, tendency to climb; 3B = indeterminate bush habit with weak mainstem and with prostrate branches, short guide, not tendency to climb). Seed color (1 = white; 2 = cream-beige; 3 = yellow; 4 = brown-maroon; 5 = pink; 6 = red; 7 = purple; 8 = black). Seed size, based on the weight of 100 seeds (1 = small, < 25 g; 2 = middle, 25-40 g; 3 = big, > 40 g)

| Line/Code | Genotype | Country of Origin | Growth type | Seed color/size | Stress response |
| --- | --- | --- | --- | --- | --- |
| MR25 | MD 23-24 x SEA 5/-(NN)C-(NN)C-25C-1C-MQ-MC | Colombia (CIAT) | Bush Bean, 2B | 2;5/2 | Drought tolerant |
| MR81 | MD 23-24 x SEA 5/-(NN)C-(NN)C-81C-1C-MQ-MC | Colombia (CIAT) | Bush Bean, 2A | 6/2 | Drought tolerant |
| MR112 | MD 23-24 x SEA 5/-(NN)C-(NN)C-112C-1C-MQ-MC | Colombia (CIAT) | Bush Bean, 2B | 6/2 | Drought tolerant |
| MR8 | MD 23-24 x SEA 5/-(NN)C-(NN)C-8C-1C-MQ-MC | Colombia (CIAT) | Bush Bean, 2A | 2;6;8/2 | Drought susceptible |
| MR109 | MD 23-24 x SEA 5/-(NN)C-(NN)C-109C-1C-MQ-MC | Colombia (CIAT) | Bush Bean, 2A | 4/2 | Drought susceptible |
| MR116 | MD 23-24 x SEA 5/-(NN)C-(NN)C-116C-1C-MQ-MC | Colombia (CIAT) | Bush Bean, 2A | 6/1 | Drought susceptible |
| BH2 | BAT 881 x G 21212/-1-1-M-M-M-M-M-M-M-M | Colombia (CIAT) | Bush Bean, 2A | 8/1 | Drought tolerant |
| BH9 | BAT 881 x G 21212/-1-1-M-M-M-M-M-M-M-M | Colombia (CIAT) | Bush Bean, 2A | 4/1 | Drought tolerant |
| BH45 | BAT 881 x G 21212/-1-1-M-M-M-M-M-M-M-M | Colombia (CIAT) | Bush Bean, 2B | 8/1 | Drought tolerant |
| BH152 | BAT 881 x G 21212/-1-1-M-M-M-M-M-M-M-M | Colombia (CIAT) | Bush Bean, 2B | 4/1 | Drought tolerant |
| BH36 | BAT 881 x G 21212/-1-1-M-M-M-M-M-M-M-M | Colombia (CIAT) | Bush Bean, 2A | 4/1 | Drought susceptible |
| BH50 | BAT 881 x G 21212/-1-1-M-M-M-M-M-M-M-M | Colombia (CIAT) | Bush Bean, 2B | 8/1 | Drought susceptible |
| SEA5 | SEA 5 | Colombia (CIAT) | Bush Bean, 2A | 2/2 | Drought tolerant |
| MD2324 | MD 23-23 |  | Bush Bean, 2A | 6/1 | Heat tolerant, moderate resistance to BGYMV |
| BAT881 | BAT 881 | Colombia (CIAT) | Bush bean, 2A | 4/1 | Elite line, drought susceptible, low<br>P sensitive |
| G21212 | G 21212 |  | Bush Bean,<br>3B | 8/2 | Excellent grain<br>filling, drought<br>susceptible |
| DOR390 | DOR 390 | Colombia<br>(CIAT) | Bush Bean,<br>2B | 8/1 | Drought<br>susceptible |
| SEN56 | SEN 56 | Colombia<br>(CIAT) | Bush Bean,<br>2A | 8/2 | Drought tolerant |
| G40001 | G 40001 |  | Climbing<br>Bean, 3B | 1/1 | Drought tolerant |
| INB841 | INB 841 | Colombia<br>(CIAT) | Bush Bean,<br>2A | 4/1 | Drought tolerant |

### Growth environment

Field experiments were carried out at the International Center for Tropical Agriculture (CIAT) in Palmira, Colombia located at 3° 29” N latitude, 76° 21”W longitude at an altitude of 965m from July – September of 2018. Characteristics of the field and rain-out shelter trial as well as soil characteristics were described previously (Polania et al., 2016). Climate data, including minimum and maximum temperature, rainfall, and pan evaporation during the field trails were collected at 15 min intervals. During the experiment, temperatures ranged between 14.1-37.2 °C, with a daytime average of 25 °C, average air relative humidity was 78%, average solar radiation was 500 watt/m^2^ and average daylight PAR of 970 µmol m–2 s–1). Total rainfall during the active crop growth period was 190 mm with potential pan evaporation 469 mm. Irrigation was maintained in the field for the control treatment, and two drought-stress treatment levels (see below) were managed under rain-out shelter conditions. The irrigated control treatment received 6 furrow irrigations (each 30 mm of water) together with two rains (55 and 30 mm) to ensure adequate soil moisture during the season. For replication, all the treatments were split into 3 separate randomized blocks each containing 4 internal columns. Each of these columns was made up of 20 plots, one for each genotype, which contained 8 individual plants. Each column had a different, random order of the plots. In total, 12 replicate plots existed per treatment. A border *Phaseolus vulgaris* genotype ‘Amadeus’ was used at each exterior edge as well as between columns. The soil is a Mollisol (fine-silty mixed, isohyperthermic Aquic Hapludoll) as described by the USDA classification system, with no major fertility problems (pH = 7.7). For a more detailed description, see Beebe et al. 2008 and Rao et al. 2017. All other information concerning this field experiment was similar to Polania et al., 2016.

### Drought treatment

Two different drought treatments were used to determine the independent effects of water stress on two different growth processes, leaflet growth rates and pod growth rates. For determining impacts to leaflet growth rates, water was withheld 10 days before leaflet growth measurements began at BBCH stage 15-17 (5-7 true leaves unfolded; 27 days after sowing) (Feller *et al.*, 1995). After 5 consecutive days of leaflet measurements, this early droughted treatment (ED) was re-watered to 80% of field capacity using 30 mm of water by sprinklers. After this re-watering, water was again withheld for the remainder of the experiment until the final harvest (69-77 days after sowing). For determining impacts to pod growth rates, a separate part of the rain-out shelter remained well-watered throughout canopy development, similarly to the control field, and only had water withheld 5 days before pod measurements began at BBCH stage 69 (end of flowering, first pods visible) (Feller *et al.*, 1995) referred to as the late drought treatment (LD). LD plants continued to have water withheld after pod measurements for the remainder of the experiment until the final harvest (69-77 days after sowing). To prevent any rainfall that might occur from disrupting either drought condition, ED and LD fields were grown under a rain-out shelter – a transparent, rolling-roof structure that was positioned over the fields whenever rain threatened (Fig. 1), otherwise remained open. Each drought treatment (ED and LD) was applied just before the specific developmental process being measured began (leaf elongation and pod elongation, respectively) in order to probe that specific process’ response to water stress while limiting impacts to other processes, such as canopy development. A third field was maintained as the well-watered (WW) treatment, adjacent to the rain-out shelter but not under it. This field was watered to field capacity every 2-3 days to maintain relatively constant volumetric water content (Fig. 2). Although the ED plants were only intended for leaflet growth rate measurements initially, surprisingly high survival allowed for pod growth measurements to be taken on these plants, as well as yield, PHI, and biomass dry weight. These data are included in some analyses.

**Fig. 1.**
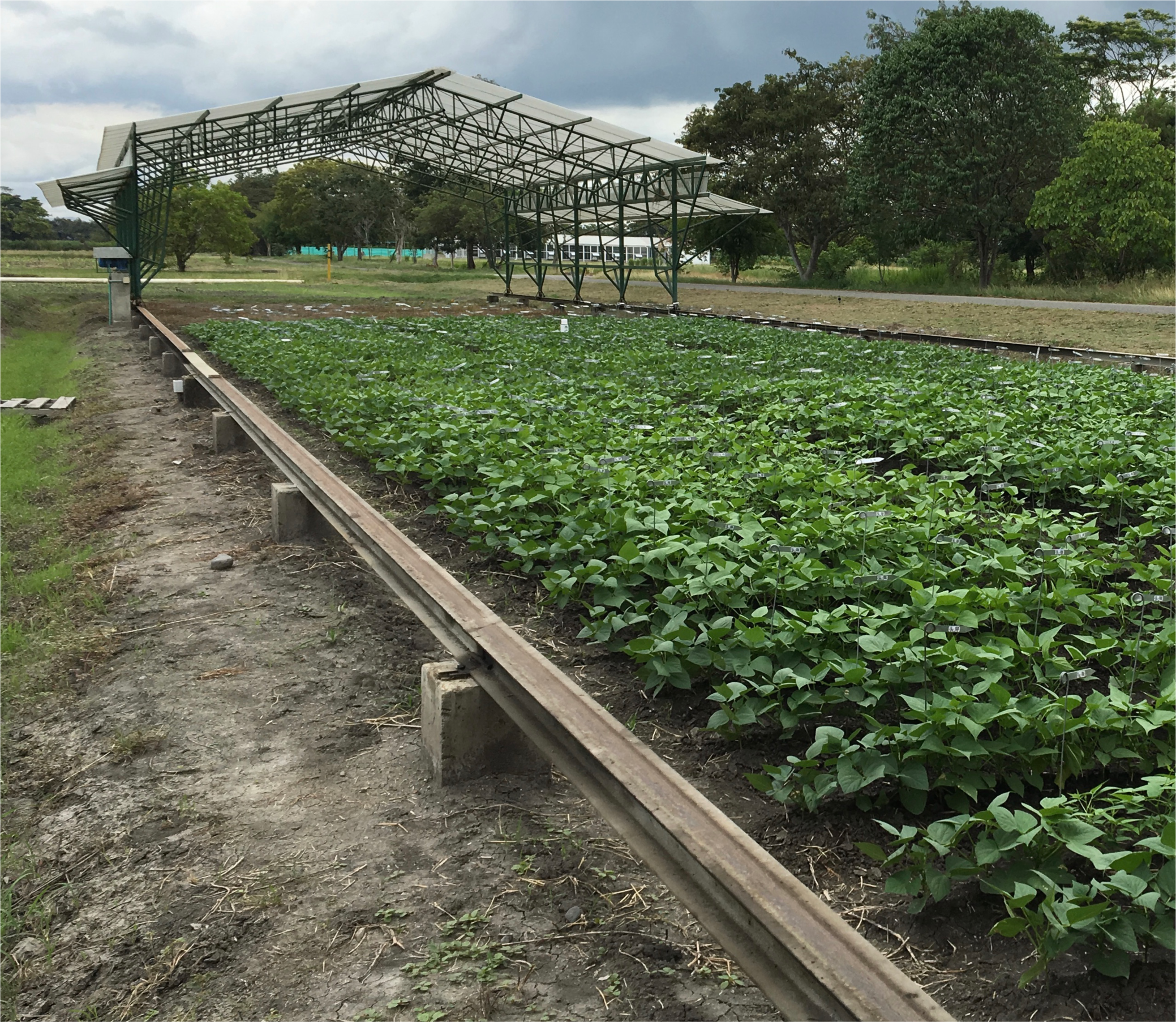
Photograph of the field and rain-out shelter, closed

**Fig. 2.**
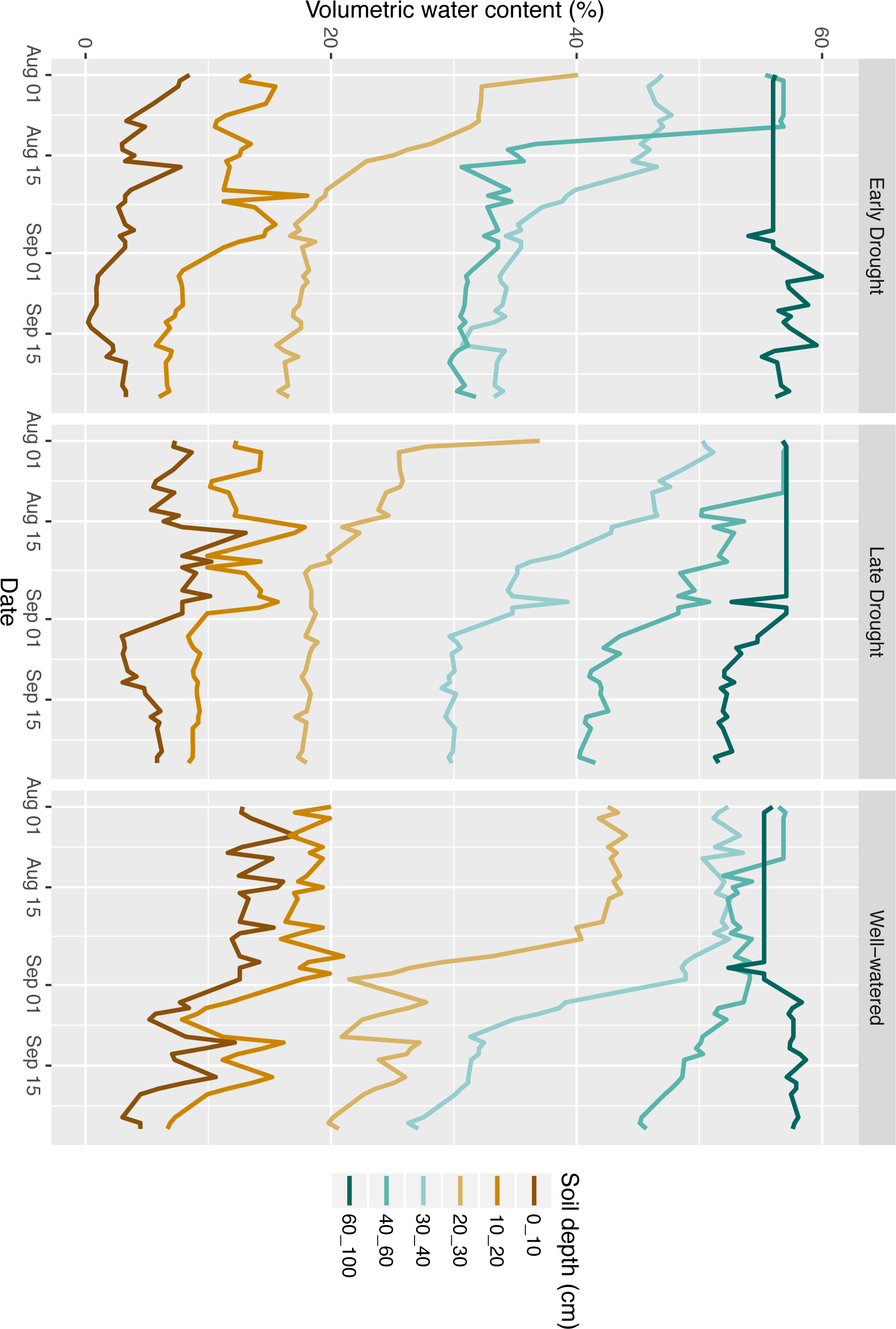
Average soil water content at different depths across the three conditions; Early Drought, Late Drought and Well-watered. Water content in depths from 0-100 cm were monitored three times daily across the three conditions, averaged, and plotted over the course of the experiment.

### Growth rates

During canopy development, when the plants were 3.5 weeks old, leaflet growth rates were measured from 3 replicate plots per treatment in both WW and ED conditions. Within each of these plots, 3 individual plants were selected, for a total of 9 replicates per treatment. For each replicate, a terminal leaflet between the length of 40-70 mm, around 30-50% fully expanded (corresponding to the start of linear elongation phase, data not shown), was tagged and blade length from base to tip was measured using a ruler and recorded. Length measurements of the same leaflet continued for a total of 5 consecutive days, each measurement taken 24 hours apart.

For pod growth rates in all three conditions, 3 individual plants were also chosen per each of three plots, and the very first and second pods which developed on each individual plant were marked. Pod lengths from base to tip were measured with a ruler daily. Measurements began when pod lengths were between 10-20 mm and continued for 3-6 consecutive days, each measurement taken 24 hours apart.

### Water potential

During the week of leaflet length measurements, water potential only for plants in ED and WW conditions was measured (since the LD condition had not yet entered its drought condition, therefore was identical to the WW treatment, data not shown). During the week of pod elongation measurements, water potential for ED, LD and WW were all measured. An individual leaf per plot was measured for each genotype, and this was repeated across two blocks. This resulted in 2 measurements per genotype per treatment. These replicates came from different plants than those marked for growth rate measurements. Near fully-expanded terminal leaflets near the top of the canopy were selected and cut at the furthest end of their petiolule from the leaflet blade using a razor blade. The leaflet was quickly put into a humid plastic bag and stored in the dark on ice until the measurement was determined, no longer than 10 minutes after cutting. Leaf water potential was taken using established protocols for a Scholander pressure chamber with a compression gasket system (model 615, PMS Instrument Co., USA). For midday measurements, leaves were collected between 1400-1600. Pre-dawn measurements were made with the same procedure between 0530-0800.

### Solute potential

After each leaflet’s water potential was measured, it was individually placed into a 2 ml Eppendorf tube and stored on ice until placed into a −20C freezer. Later, solute potential was determined using established protocols with a vapor pressure osmometer (model 5100B, Wescor Inc., USA). Samples were thawed for 20 minutes, exposed to remove condensation, and slightly pressed between two microscope glass slides to release sap which was then measured. Three measurements were taken per sample and averaged.

### Yield, PHI, Biomass

Upon physiological maturity, 3 consecutive plants were destructively harvested from each of 6-13 replicate plots. All dry seeds from these 3 plants were weighed together then divided by 3 to give dry seed yield per plant. Whole pod dry weight (including seeds) was also determined for the same 3 plants together in each plot and PHI was calculated per each replicate plot using equation 1.

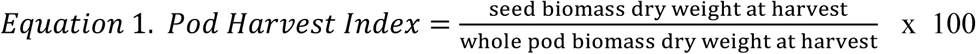

Whole canopy (above-ground) dry biomass was averaged for the same 3 plants weighed together from each plot.

### Statistical analysis

Linear regression correlations were made between traits which were then tested for statistical significance via the student’s t-test using Microsoft Excel (one-tailed, unpaired) alpha = 0.05 level of significance. Graphs represent mean values ± standard deviation.

When describing results, absolute values and impacts are reported, where absolute values refer to actual recorded values, whereas impacts refers to a calculated percent decrease between the droughted and WW values for a trait (equation 2). The genotypes that decreased by a larger percentage were considered to be more impacted and drought sensitive.

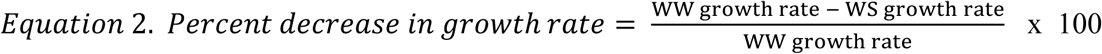

## RESULTS

### Leaf water potential and solute potential

To quantify internal water status within the plants, leaf water potential was measured predawn and midday. Measurements were taken over multiple days and values from 2 or 3 days of measurements were averaged by condition and graphed by week (Fig. 3A). As intended, when all genotypes were averaged together, water potential values were significantly lower in water-stressed conditions compared with WW, showing the drought condition resulted in water deficit compared to WW. This was true for all pre-dawn and midday WW to water-stressed comparisons.

**Fig. 3.**
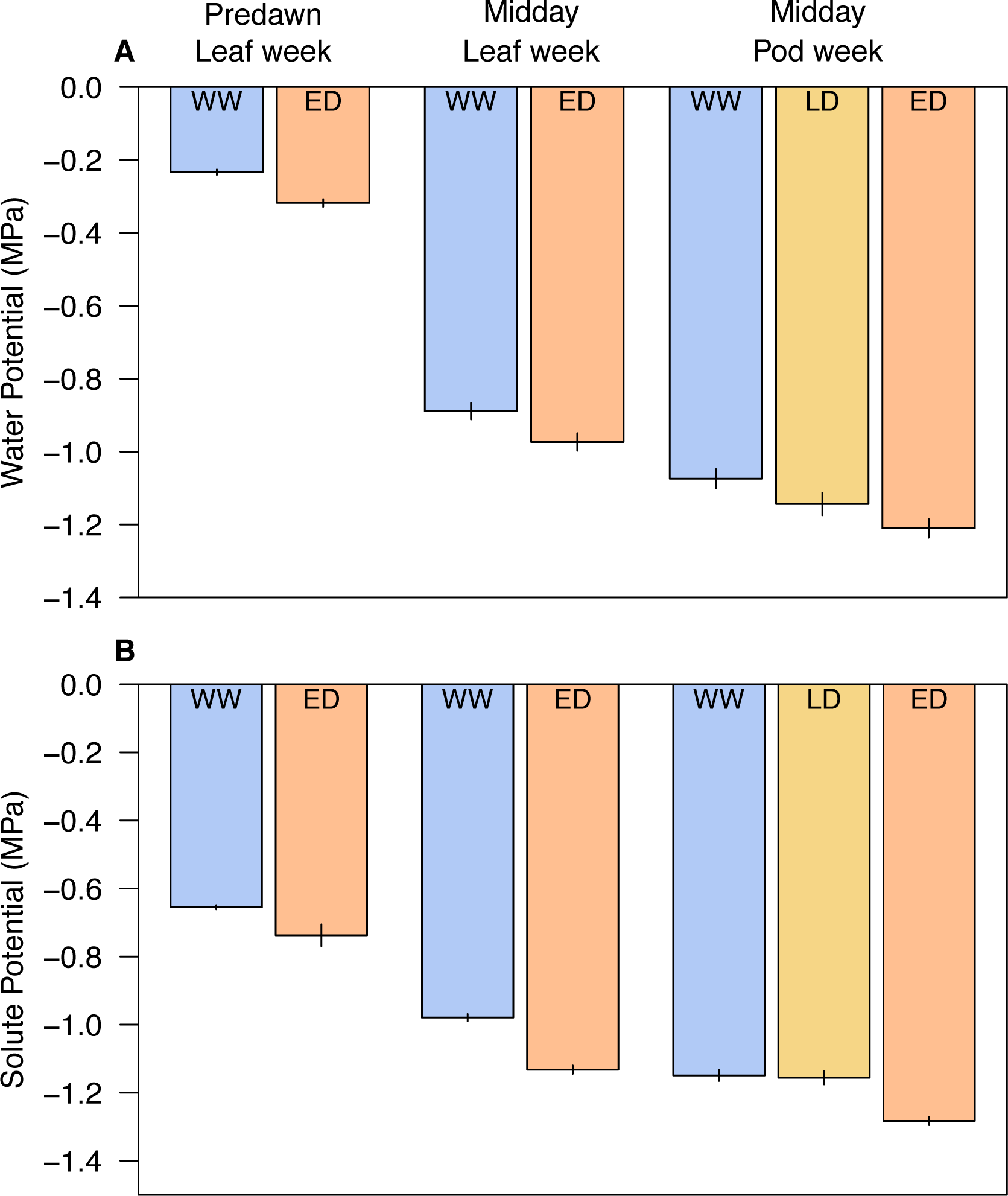
Average leaf water potential (A) and solute potential (B) at predawn or midday during the two weeks of growth measurements. ‘Leaf week’ refers to when leaflet growth rate measurements were taken, ‘Pod week’ when pod growth rates were taken. All bars represent the average across all 20 genotypes, using 2 or 3 measurements per genotype. Error bars show standard error.

When each genotype was looked at individually, grouped by treatment and time point, difference among genotypes within a condition/timepoint ranged from −0.31 MPa (for WW/week of leaflet elongation, which had the most similar values between genotypes) to −0.54 MPa (for LD/week of pod elongation, which had the most different values between genotypes). Even though variation existed in leaf water potential between genotypes, leaf water potential value by genotype did not correlate with the other physiological or agronomic traits measured for that genotype, such as leaflet and pod growth rates, biomass and PHI (PHI shown). This suggests that maintaining higher (closer to zero) leaf water potential values did not result in faster growing leaflets or pods, nor with higher seed yield, PHI or canopy biomass. Among the individual genotypes, there was also a range in how impacted they were by drought. When values were compared between ED and WW during the week of leaflet elongation, some genotypes had little to no decrease in water potential, while others decreased by close to 50% of the WW values. However, as with the absolute water potential values, genotypes whose water potential was most impacted by drought were not the same genotypes whose growth rates, PHI, yield, or biomass were most impacted (data not shown).

Solute potential trends were similar to water potential, with values typically decreasing between droughted and WW (ED for leaflet measurement week, LD for pod measurement week) (Fig. 3B). One notable difference is that solute potential of LD vs WW during the week of pod measurements were not different from each other. Lastly, as with water potential values, solute potential values and impacts to solute potential values did not correlate with growth or other agronomic traits nor the impacts to growth or agronomic traits, respectively.

### Leaflet and pod growth rates

As expected, most leaflet growth rates (measured in the ED condition) were significantly impacted by drought, however, fewer pod growth rates (measured in the LD condition) were significantly impacted. Leaflet growth rates always decreased between ED and WW, but the amount of decrease varied widely by genotype, ranging from 3% to 46% (Fig. 4A). All decreases of 20% or more were significant, which was the case for 16 of the 20 genotypes. Pod growth rates also typically decreased between LD and WW treatments, by around 2% to 45%, although for three genotypes (MR116, INB841, and BH50), pod growth rates increased under LD (Fig 4B). Of those three, only increases in BH50 were significant. Impacts to pod growth rates were only significant for 7 of the 20 genotypes. Yet, since both leaflet and pod growth rates decreased by similar extents, we hypothesized that genotypes whose leaflet growth rates were most impacted would also have the most impacted pod growth rates. This would have suggested that drought impacts growth processes in different tissues in a common or conserved way. However, the degree to which these different tissue types were impacted was not consistent across genotypes; genotypes whose leaflet growth rates decreased the most under drought did not have pods whose growth rates decreased most. Instead, these two growth rates seemed to be independently impacted by drought stress within a genotype (Fig. 4 inset).

**Fig. 4.**
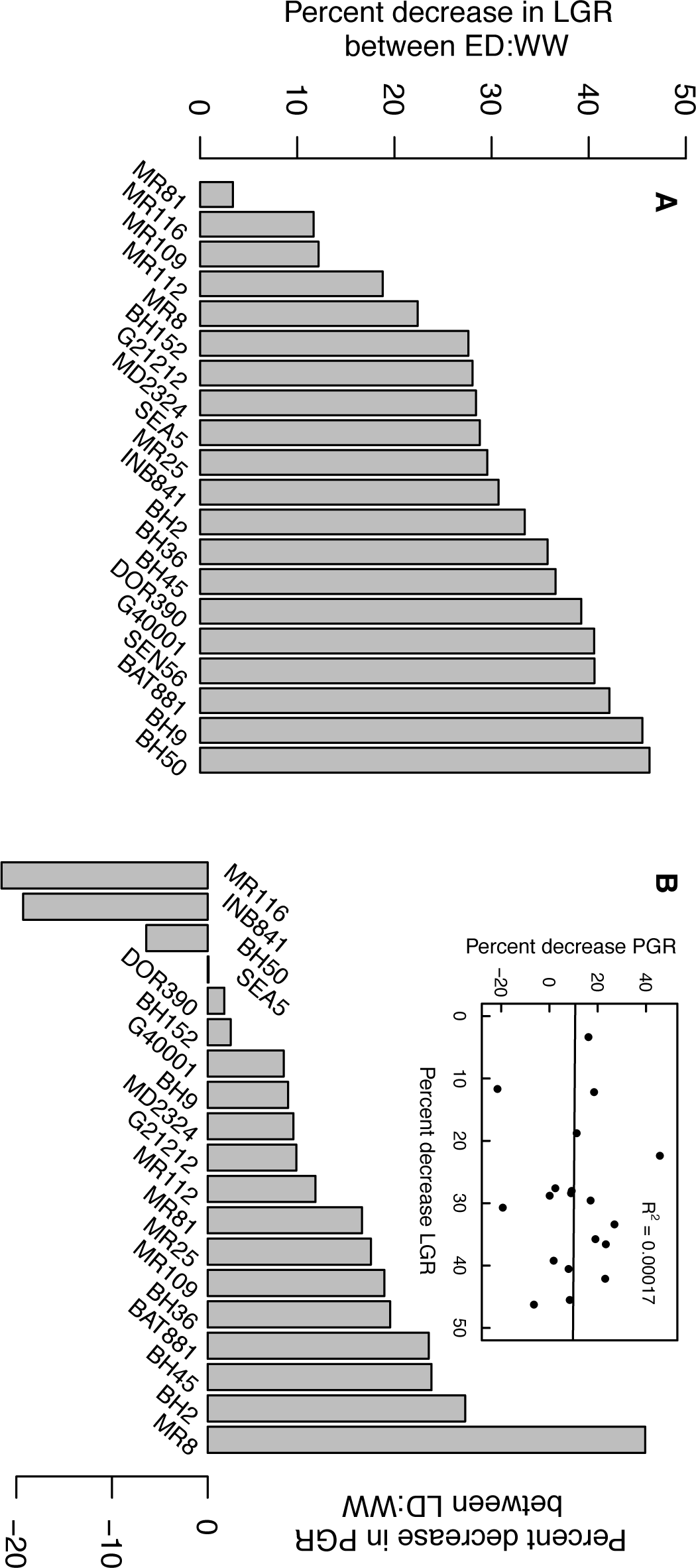
Impacts of drought on leaflet growth rates (A) and pod growth rates (B) by genotype. Impact is shown as percentage decrease in rate between Well-watered (WW) and droughted; early drought (ED) for leaflet growth rate impacts and late drought (LD) for pod growth rate impacts. A negative percent decrease, as occurred for four genotypes in B, indicates an increase in rate under LD compared to WW. Inset shows correlation between impact of drought on leaflet and pod growth rates.

### Growth rates and seed yield

Leaflet growth rates had a weak but significant correlation with yield under WW. However, neither leaflet growth rate nor pod growth rates were correlated with yield under WS conditions, nor was there correlation between pod growth rates and yield under WW conditions (Fig. 5A and B).). Instead, the impact of drought on leaflet growth rate was better correlated with the impact of drought on yield. Specifically, we found that the genotypes whose leaflet growth rates were most impacted by drought (compared between ED:WW) also tended to have seed yield highly impacted (compared between LD:WW). This was true among the 19 *P. vulgaris* genotypes (Fig. 5C). Interesting, while *P. acutifolius* had very low impact on yield under WS as expected, leaf growth rate was highly impacted, showing that leaf growth and yield impacts in this species are more decoupled. Unlike leaflet growth rates, pod growth rate sensitivity to drought did not correlate with yield decreases under drought stress (Fig. 5D).

**Fig. 5.**
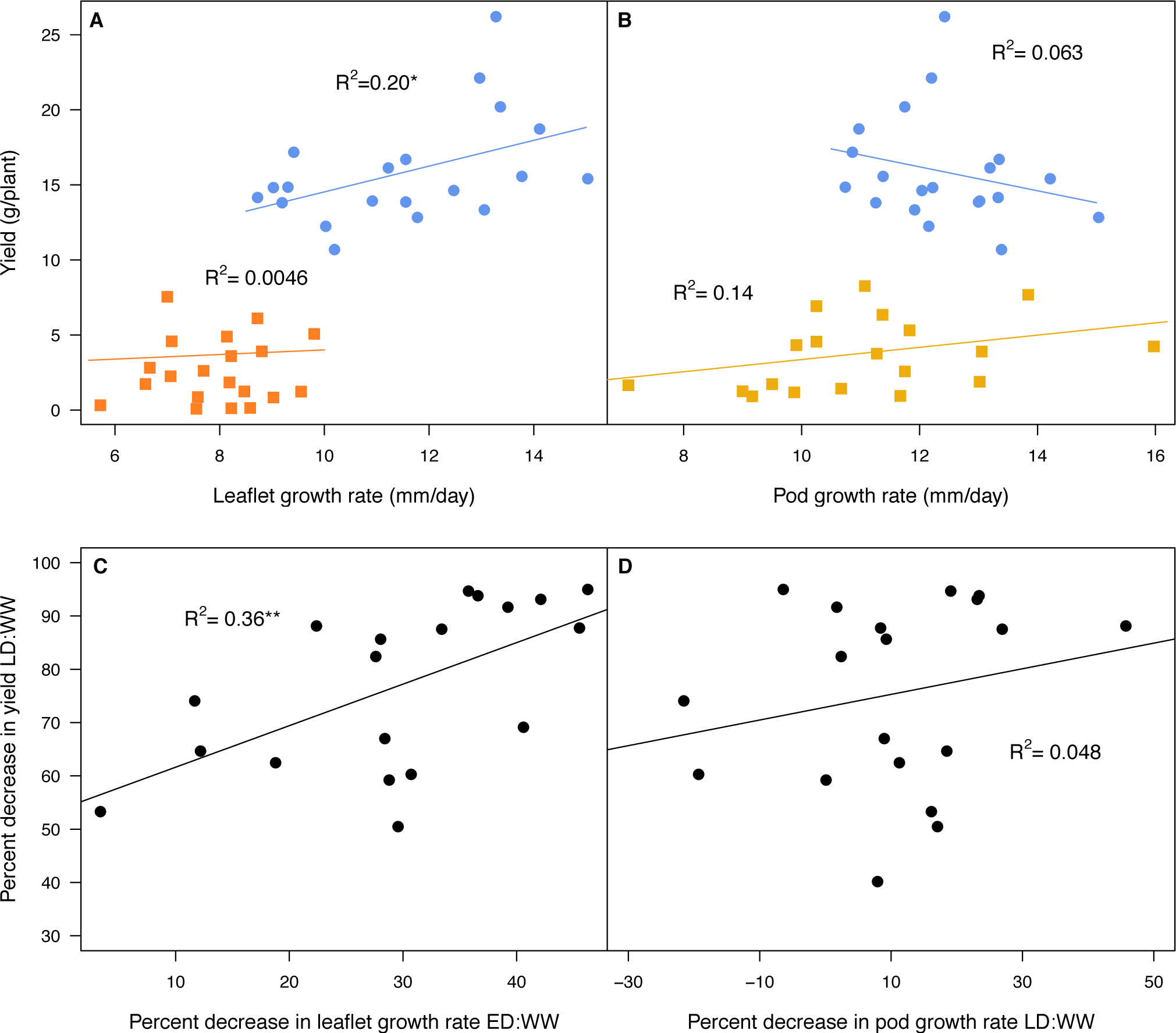
Correlations between yield and growth rates. In A, correlations were tested between average WW leaflet growth rate and WW yield (blue) and ED average leaflet growth rates and LD yield (orange). B shows the same but for pod growth rates and yield under LD (yellow). In A and B, circles denote WW and squares represent water stressed. C and D tested correlations between impacts to yield (percent decreases between LD and WW) and (C) impacts to leaflet growth rates (percent decrease between ED and WW) or (D) impacts to pod growth rates (between LD and WW). Note that in A and C, ED leaflet growth was compared to LD yield, whereas in B and D, LD pod growth was compared to LD yield.

Furthermore, genotypes with the highest leaflet growth rates under WW conditions (BH50, BH9, and BAT881) were the most impacted under ED (Fig. 6). This result, in combination with the fact that these genotypes also experienced strong decreases in yield under LD (95%, 88% and 93%, respectively), suggests that genotypes with the fastest LGR under WW conditions are at high risk of negative impact to leaflet growth rates and seed yield under drought. However, genotypes with the fastest growing pods under WW conditions did not have pod growth rates that were most impacted under LD (data not shown – R^2^=0.08).

**Fig. 6.**
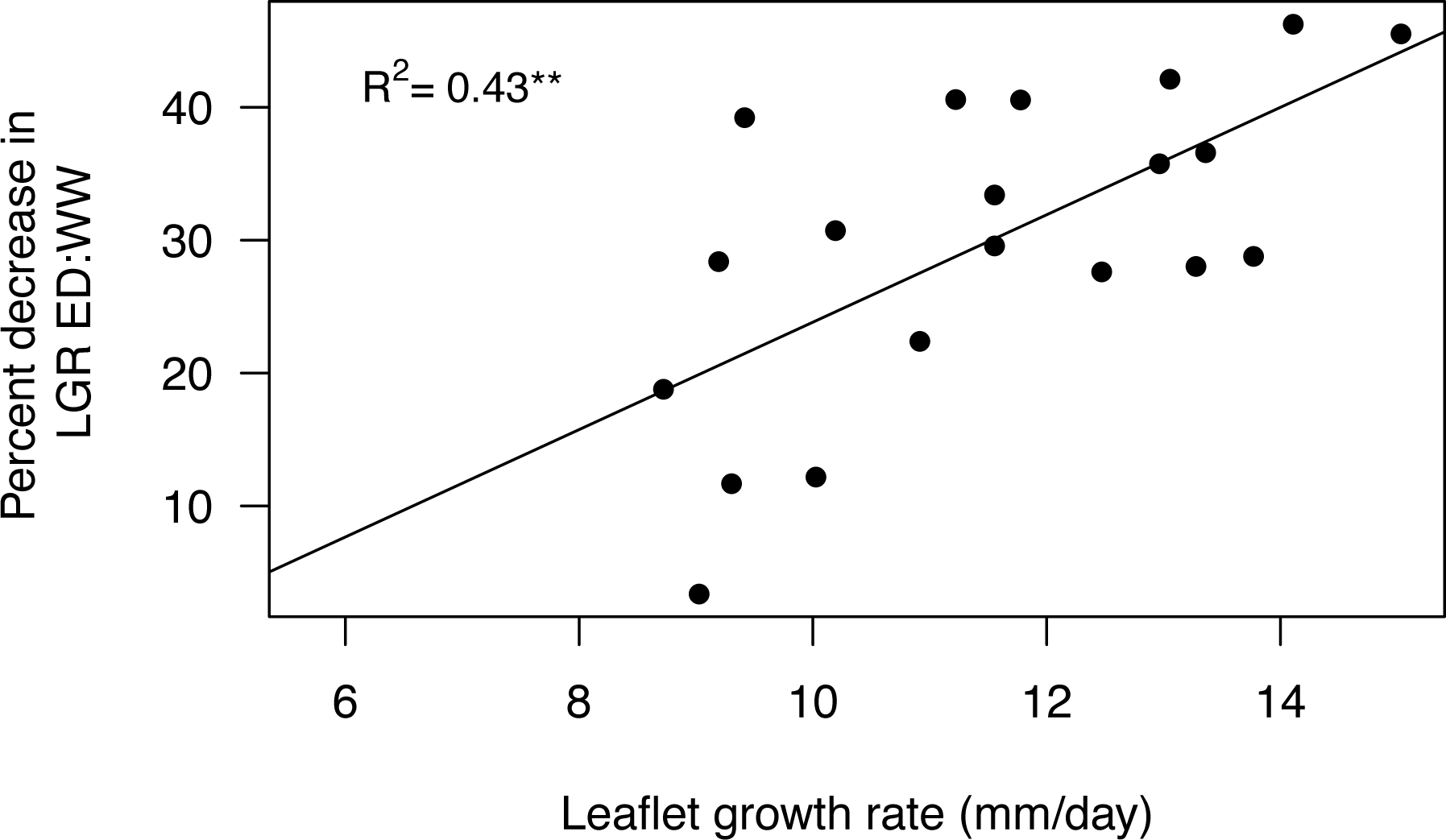
Correlation between WW leaflet growth rates and impacts to leaflet growth rates

### PHI

Under WW, average PHI values ranged from 0.74 to 0.83, with few significant differences among genotypes. However, under LD, average PHI values ranged from 0.47 to 0.78 and under ED even wider, from 0.24 to 0.77 (Fig. 7). While differences amongst WW genotypes were small and rarely significant, larger differences existed amongst both the ED and the LD genotypes individually, as well as differences between each drought condition and WW for most genotypes. Although significant decreases between WW to ED and LD existed for most genotypes, the amount of difference between droughted and WW values varied widely by genotype from 0.02 to 0.31 under ED and 0.03 to 0.54 under LD. The genotype with not only one of the highest PHI values under WW but also the least impacts to PHI under either drought condition was G40001, *P. acutifolius*, which was included because it was known to maintain high PHI under drought.

**Fig. 7.**
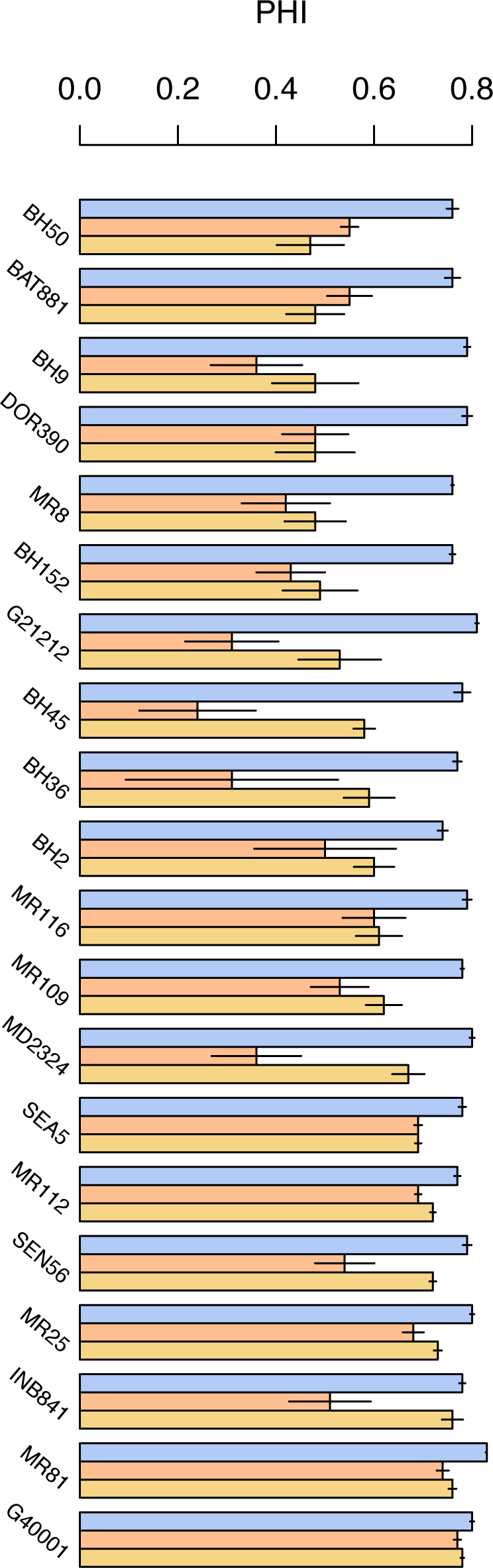
PHI values for all 20 genotypes under WW (blue), ED (orange), and LD (yellow) conditions. Genotypes have been ordered in ascending order based on LD PHI values. Error bars show standard error.

When PHI values were compared against LGR and PGR, no significant correlations were found in either WW or WS conditions, nor impacts to these values between well-watered to water stressed (data not shown).

### PHI and yield

Studies from CIAT consistently show PHI correlates more strongly with seed yield under drought and well-watered conditions than biomass, with these correlations typically strongest in droughted plants over those growing in WW conditions (Rao *et al.*, 2017). In our study, PHI did not correlate with yield under WW conditions, however, under water deficit the relationship between PHI and yield was high, as expected, with significant positive correlation under LD conditions (Fig. 8A). Likewise, decreased in PHI and yield under LD have significant positive correlation (Fig. 8B).

**Fig. 8.**
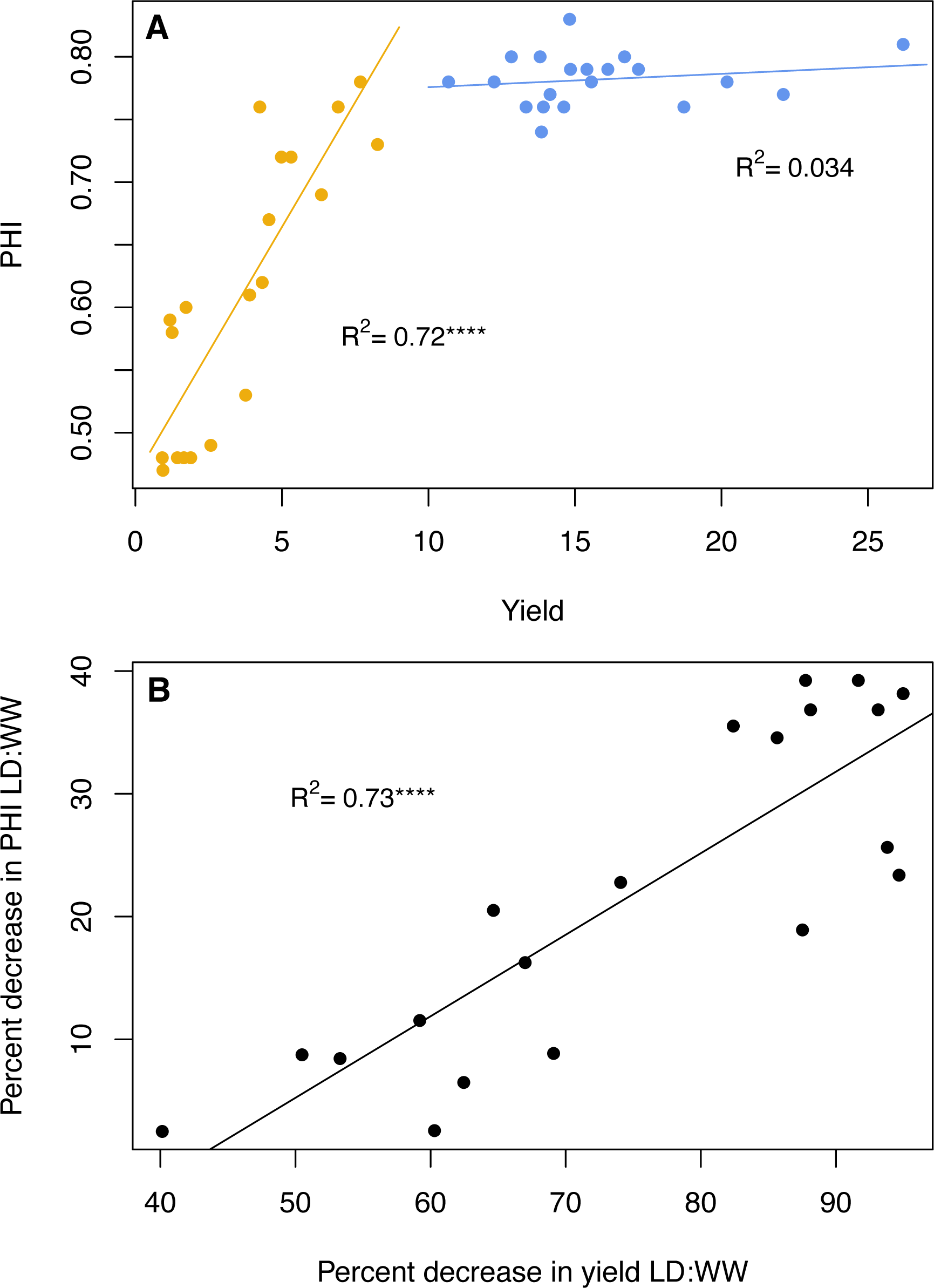
Correlations between yield and PHI. In A, correlations were tested between absolute yield and PHI under WW (blue) and LD (yellow) for the 20 genotypes. B shows correlations between impacts to these two traits comparing between LD to WW.

### Canopy biomass and seed yield

While canopy biomass was correlated with seed yield under LD conditions (data not shown – R^2^ = 0.59), the correlations between the impacts on these two traits were even higher. Specifically, impact on canopy biomass under LD correlated more strongly with seed yield than any other trait (Fig. 9); those plants with the largest decreases in canopy biomass also had the largest decreases in seed yield. This suggests that while biomass itself is a large contributor to seed yield potential, the relative sensitivity of a plant’s biomass accumulation to drought was a better predictor of yield over absolute values of biomass itself.

**Fig. 9.**
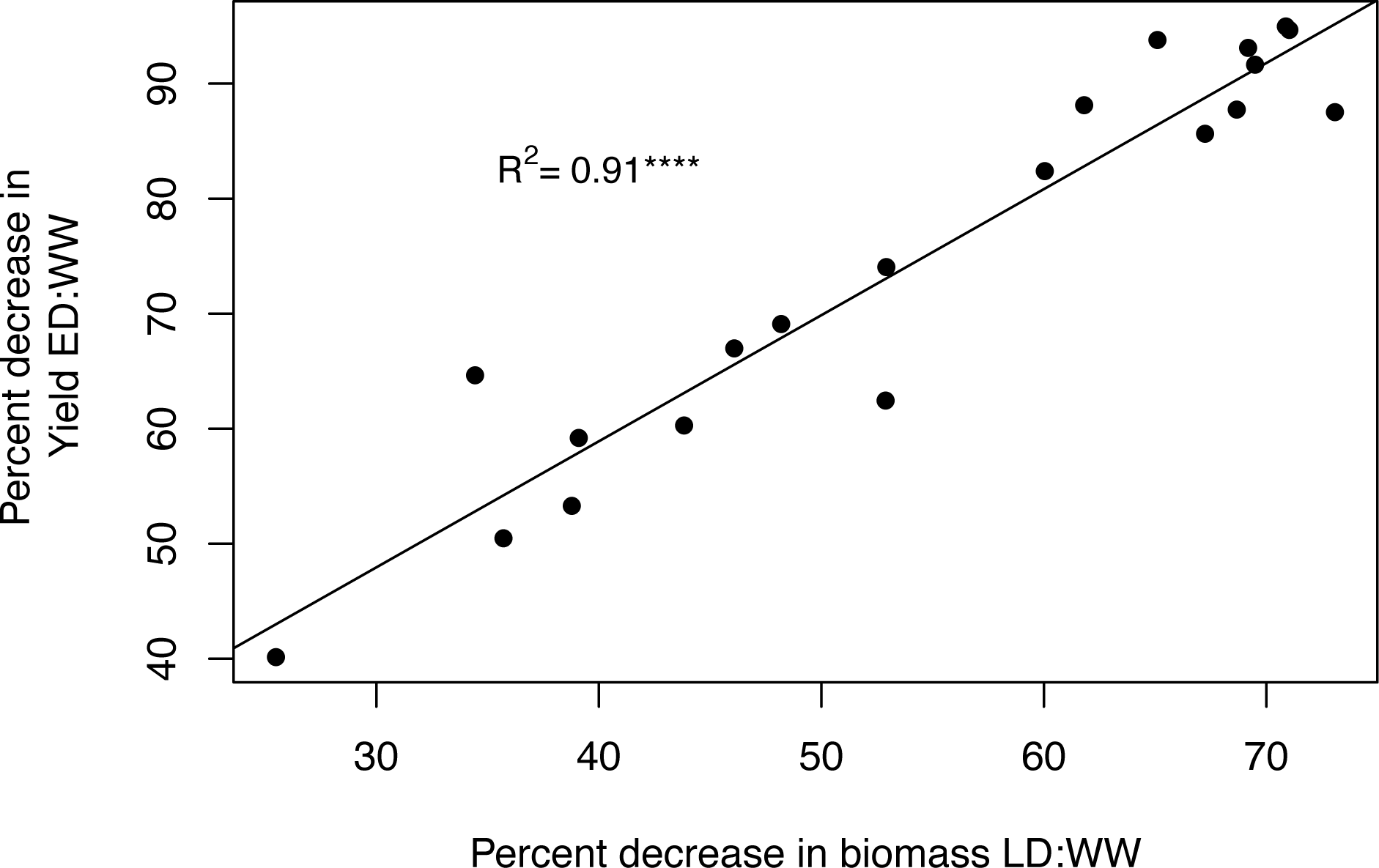
Correlation between impacts to yield and biomass between LD and WW conditions.

## DISCUSSION

As intended, both ED and LD water-stress conditions resulted in water deficits within the plants when compared to WW. Surprisingly, leaf water status did not seem to play a direct role in limiting growth and yield, since neither leaf water potential nor solute potential values correlated with growth rates, canopy biomass, PHI or yield. Nor were genotypes whose water or solute potentials were the most impacted by drought the same genotypes whose growth rates, canopy biomass or PHI were the most impacted. This suggests that leaflet and pod elongation, biomass accumulation, and resource partitioning are not limited directly by low leaf water potential, since genotypes with the lowest leaf water potential did not have lowest values in the above-mentioned traits.

Impacts of drought on leaflet and pod growth rates within each genotype were uncoupled, meaning genotypes whose leaflet growth rates decreased the most (under ED – Fig. 4A) were not the same as those whose pod growth rates decreased most (under LD – Fig. 4B). Although we had predicted drought responses across tissue types would be similar, due to conserved drought-sensitivity mechanisms, we found leaflet growth rate impacts could not be used as an indicator of how pod growth rates would be impacted (Fig. 4 inset). This could be due to the fact that under drought, tissues types become water stressed at different rates, with pods and seeds being the last impacted (Westgate and Grant, 1989). Pods may be buffered from this stress because of the critical role they play in survival which may allow growth processes to be less affected in pods. Meanwhile in leaves, which are experiencing stronger water deficit, drought signals such as abscisic acid (ABA) may be hindering growth. And while ABA can limit photosynthesis, through causing stomata to close, ABA accumulation also signals inhibition of wall loosening and cell growth in growing leaves, which could separate this phenomenon from substrate limitation (Davies and Van Volkenburgh, 1983). Drought signals may still affect leaflet and pod tissues similarly, but pods in our study may not have experienced the same level of stress as leaves. Future work could dissect this with measurements of water potential in these different tissues as well as through quantification of stress indicators within them, including ABA levels or dehydrin accumulation etc.

We predicted that leaflet and pod growth rates would correlate with seed yield and partitioning efficiency (PHI). We found that while leaflet and pod growth rates themselves did not correlate with partitioning or yield under water deficit, the sensitivity of leaflet growth rate to drought did predict the sensitivity of yield to drought. In other words, genotypes whose leaflet growth rates were most impacted by drought in comparison to control had yields most impacted as well. We had predicted that genotypes which were able to maintain higher LGR under drought may do so via maintenance of high sink strength in leaves. This in turn could allow them to maintain higher pod growth rates and seed filling, which would ultimately lead to higher yield. This hypothesis was based on published results where terminal drought tolerance was explained by higher efficiency of carbon mobilization from leaves to pods and seeds (Cuellar-Ortiz *et al.*, 2008; Rosales *et al.*, 2012). This does not appear to be true when looking at absolute values of growth rates and yield, as some high yielders had lower leaflet and pod growth rates (Fig. 5A & 5B). Yet it does appear true in terms of sink strength sensitivity, where impacts to leaflet growth and yield seem to be linked by a conserved mechanism affecting processes related to sink strength (Fig. 5C). These data suggest that relative LGR sensitivity to drought may act as a good predictor of overall drought resistance.

Unsurprisingly, given that LGR and PGR within a genotype were not similarly impacted by drought, PGR impacts under LD were not a good predictor of yield (Fig. 5D), which is the opposite of the result under LGR. This could again be because the pod is buffered from water stress, separating its response from other sink tissues. However, for the ED treatment, absolute PGR and yield did have a significantly positive correlation. Since we only saw this to be true under ED, we believe this could mean that more severe stress (especially when it leads to large impacts to biomass, as it did in the ED treatment) impacts PGR due to lack of resources, whereas under the LD treatment, genotypes that slowed most did so due to a stronger response to drought signaling or status, rather than a lack of substrate.

Unlike leaflet and pod growth rates, canopy biomass values did correlate strongly with seed yield under both WW and LD conditions. Yet, stronger still were correlations between impacts to biomass and impacts to yield under LD (Fig. 9). This again supports the above-mentioned hypothesis that genotype sensitivity to drought may be its strongest predictor of yield under drought. Yet, particularly under LD conditions, genotypes in this study which achieved a similar biomass displayed a range of yields, with some genotypes differing even up to 80%. Therefore, while the correlations show that higher biomass is necessary to achieving higher seed yield under drought stress, as has been shown previously, (Polania et al. 2017; I. M. Rao et al. 2017), reductions in canopy biomass alone does not fully answer the question as to what results in reduction of yield under drought.

Therefore, based on previous findings that PHI is the best predictor of yield, we assessed whether PHI values help to account for differences in yield when biomass could not. First, we tested correlations between PHI and yield under the three conditions. We found that PHI did not correlate with yield under WW conditions. However, we did find correlations under ED conditions and even higher under LD conditions (Fig. 8A). We also found that impacts to yield and PHI between LD and WW correlated significantly (Fig. 8B) Beyond testing how PHI alone related to yield, we wanted to know whether differences in PHI could be used to better understand how genotypes can acquire the same canopy biomass yet achieve different yields. Indeed, we found they could. For example, under LD, average canopy biomass ranged from 20-55g per plant, depending on genotype. As mentioned above, within that range, genotypes whose canopy biomass were the same could vary by 80% in their yield. Specifically, genotypes MR81 and BH45 had very similar high average canopy biomass under LD – around 43g. Yet while MR81 yielded second highest of all the genotypes under LD, with 6.72g dry seed weight, BH45 was on the lower end of the yield spectrum with only 1.25g. If 6.72g is considered 100% of the potential yield possible for this canopy biomass, a yield of 1.25g is only 19% of that potential. When we paired this finding with PHI values for these genotypes, the differences in yield between the two could be explained by differences in their partitioning efficiency, with MR81 maintaining a high PHI of 0.76 while BH45 had a much-reduced value of 0.58 (Fig. 7).

Our results help to tease apart systemic and tissue-specific responses to drought and to understand how impacts to different growth or partitioning processes under drought relate to yield. While our results did not support the prediction that leaflet and pod growth rates predict seed yield, our findings together suggest that inherent differences in partitioning efficiencies and drought sensitivity may underlie a mechanism for drought resistance shared across stages of plant development. Future work will explore physiological mechanisms regulating leaflet growth and PHI, and how they are impacted under drought, to better understand this mechanism. While this study does not indicate clear physiological mechanisms to answer the question of what allows for filling of seeds in tolerant lines under drought, there are valuable agricultural insights. Plants with fastest leaflet growth had highest negative impacts on their leaflet growth under ED. Those genotypes whose leaflet growth rates were most impacted also had yields most impacted. Gaining a deeper understanding of how drought sensitivity impacts a plant’s whole life cycle (from canopy development, to flowering and pod production, to seed filling and maturation) may allow for larger gains in efficiency in yield.

Many crops whose yield has improved during the green revolution did so not by increasing their total production, but instead by partitioning a greater amount of resources to yield (Wardlaw, 1990). For example, rice and wheat yield went from having 30% of total biomass in yield at maturity to 50% in the 1960s (Khush, 1999). *Phaseolus vulgaris* yield partitioning efficiency on the other hand, has yet to see similar improvements as common bean maintains the ancestral trait of delayed seed production under drought (Beebe *et al.*, 2008, 2013). The observation that partitioning appears to limit yield under drought might fuel work to attain further improvements to partitioning in *Phaseolus vulgaris*. Gaining a deeper understanding of how partitioning is impacted by drought may allow for larger genetic gains in efficiency in this trait.

This research has the potential to increase basic understanding of plant physiology and to improve crop yields. Climate change (particularly change in precipitation distribution) is affecting soil fertility and soil water availability. In Colombia, 80% of *Phaseolus vulgaris* smallholders’ production is intended for national consumption. However, with much of the production under rainfed conditions on smallholder farms, yields are increasingly threatened. Over the past decades, the *Phaseolus vulgaris* breeding program in CIAT succeeded in identifying genotypes with increased resistance to drought, yet the mechanisms which contribute to this resistance aren’t fully understood. By identifying physiological traits to assist in developing improved *Phaseolus vulgaris* cultivars, this work attempts to contribute to more stable yields and food security. Discoveries on sink strength in common bean may also help to uncover common mechanisms shared by other crops, increasing the impact of our findings.

## ACKNOWLEDGMENTS

We thank Dr. Idupulapati Rao, Dr. Steve Beebe, and Dr. Maria Reguera for invaluable discussions and input leading to this project. Our thanks also to Jose Polania and to the field team at CIAT for their work and dedication. We wish to recognize the USAID project “Bean Improvement for Tropical Producers and Consumers: Tomorrow and Beyond” and the TL-3 Project “Improving Livelihoods for Smallholder Farmers: Enhanced Grain Legume Productivity and Production in Sub-Saharan Africa and South Asia” for funding, as well as the Department of Biology at the University of Washington for travel funds making this research possible.

## ABBREVIATIONS

ABA: abscisic acid
CIAT: International Center for Tropical Agriculture
ED: early drought
LD: late drought
LGR: leaflet growth rate
PGR: pod growth rate
PHI: pod harvest index
RIL: recombinant inbred line
WW: well-watered

